# Altered infective competence of the human gut microbiome in COVID-19

**DOI:** 10.1101/2022.10.20.512999

**Authors:** Laura de Nies, Valentina Galata, Camille Martin-Gallausiaux, Milena Despotovic, Susheel Bhanu Busi, Chantal J. Snoeck, Lea Delacour, Deepthi Poornima Budagavi, Cédric Christian Laczny, Janine Habier, Paula-Cristina Lupu, Rashi Halder, Joëlle V. Fritz, Taina Marques, Estelle Sandt, Marc Paul O’Sullivan, Soumyabrata Ghosh, Venkata Satagopam, CON-VINCE Consortium, Rejko Krüger, Guy Fagherazzi, Markus Ollert, Feng Q. Hefeng, Patrick May, Paul Wilmes

## Abstract

**Objectives:** Infections with SARS-CoV-2 have a pronounced impact on the gastrointestinal tract and its resident microbiome. Clear differences between severe cases of infection and healthy individuals have been reported, including the loss of commensal taxa. We aimed to understand if microbiome alterations including functional shifts are unique to severe cases or a common effect of COVID-19.

**Design:** We used high-resolution systematic multi-omic analyses to profile the gut microbiome in asymptomatic-to-moderate COVID-19 individuals compared to a control group.

**Results:** We found a striking increase in the overall abundance and expression of both virulence factors and antimicrobial resistance genes in COVID-19. Importantly, these genes are encoded and expressed by commensal taxa from families such as Acidaminococcaceae and Erysipelatoclostridiaceae, which we found to be enriched in COVID-19 positive individuals. We also found an enrichment in the expression of a betaherpesvirus and rotavirus C genes in COVID-19 positive individuals compared to healthy controls.

**Conclusion:** Our analyses identified an altered and increased infective competence of the gut microbiome in COVID-19 patients.

## Introduction

Coronavirus disease 2019 (COVID-19), which is caused by the severe acute respiratory syndrome coronavirus 2 (SARS-CoV-2), was declared a global pandemic by the World Health Organisation (WHO). COVID-19 exhibits a high degree of clinical heterogeneity, ranging from asymptomatic to severe disease, and may be accompanied by a poor outcome and a relatively high mortality rate [1]. As of 17 October 2022, more than 621 million confirmed SARS-CoV-2 infections and 6.5 million COVID-19-related deaths have been reported [2]. Although COVID-19 is primarily considered a respiratory disease, it clinically often presents with general (fever, myalgia and/or fatigue) and respiratory symptoms (cough and/or dyspnea). Moreover, an emergence of new variants has led to the more frequent presentation of gastrointestinal symptoms (appetite loss, nausea, vomiting and diarrhoea) [3], indicating a potential involvement of the gastrointestinal tract in COVID-19. More specifically, SARS-CoV-2 has been shown to be able to infect and replicate in enterocytes *in vitro* [4]. In fact, viral RNA can be detected in faecal samples even after resolution of respiratory symptoms [5]. Additionally, SARS-CoV-2 infections are associated with alterations to the gut microbiome composition that persist for at least six months after the initial infection [6]. Thus, an imbalance in the gut microbiome can be linked to disease severity and increased concentrations of inflammatory markers, as well as an increased post-COVID-19 risk, understood as a wide range of symptoms persisting four or more weeks after the initial SARS-CoV-2 infection [6–8].

Stable ecosystems are important for colonisation resistance to pathogens [9,10]. As such, host and SARS-CoV-2-mediated immune dysregulation and dysbiosis may predispose patients to co-infections or secondary infections of the respiratory and gastrointestinal tracts. In addition, co-infecting microorganisms may alter the intensity of the host immune response [11], thus significantly influencing severity and outcome of the disease. For instance, co-infections with viruses (rhinovirus/enterovirus, respiratory syncytial virus, influenza virus, non–SARS-CoV-2 Coronavirus) [12,13], bacteria (Mycoplasma pneumoniae, Pseudomonas aeruginosa, Haemophilus influenzae, Klebsiella pneumoniae, Streptococcus pneumoniae, Staphylococcus aureus) [14,15], or fungi (Candida spp., Aspergillus spp.) [16] have been described among SARS-CoV-2 positive cases in different study set-ups. In particular, bacterial co-infections in hospitalised and intensive care unit patients with COVID-19 are associated with prolonged ventilation and an increased mortality rate [14,17–19]. Furthermore, hospital-acquired infections with multi drug resistant (MDR) pathogens are also linked with increased mortality in COVID-19 patients [20,21]. These reports collectively suggest a clear shift in COVID-19 patients with respect to an increased abundance of pathogens and potential for harm. Moreover, these shifts may further manifest themselves in relation to the infective competence, i.e. the propensity for virulence and increased antibiotic resistance, in the gut microbiome as a consequence of an increased capacity to cause infections.

Major factors that contribute to the success of some of the pathogens highlighted above are virulence factors. Virulence factors including cell-surface structures, adhesins, siderophores, endo- and exotoxins enable pathogens to undergo quick adaptive shifts, invade and colonise host niches, as well as evade innate and adaptive immune mechanisms of the host, resulting in inflammation and clinical manifestations of the disease. Another factor facilitating colonisation of pathogens, through prevention of effective treatment, is antimicrobial resistance (AMR). Even though AMR is an ancient and natural phenomenon [22], it is usually linked to the human influence on the environment and the use of antibiotics. Overuse of antibiotics is hypothesised to also contribute to the broader problem of antimicrobial resistance [23]. Moreover, although not a virulence factor by itself, AMR shares common characteristics with virulence factors [24]. Specifically, AMR and virulence factors: 1) are necessary for the survival of pathogens under unfavourable conditions [25]; 2) can be transmitted between species by horizontal gene transfer [26]; and 3) both processes make use of similar systems, e.g. cell wall alterations, efflux pumps, porins and two-component systems to activate or repress expression of various genes [24,27,28]. Thus, in response to host defence mechanisms and environmental challenges, communities of microorganisms, i.e. microbiomes, may alter their “*infective competence”.* The *infective competence* is defined as the ability of microorganisms to constantly adapt and evolve, utilising virulence factors and antimicrobial resistance mechanisms, resulting in increased survival, invasion, or growth. Importantly, the combination of host-driven factors, i.e. immune system-mediated effects and antimicrobial peptides, and unfavourable gastrointestinal conditions, e.g. low pH, disruption of the mucus layer, niche competition with other taxa, may confer transiently a selective advantage to a pathogenic lifestyle [29–32]. This may be reflected in the entire gut microbiome, possibly altering the *infective competence* of the endogenous taxa and subsequently giving rise to pathobiont-dominated communities.

Here, we addressed questions pertaining to the effect of SARS-CoV-2 infection on the endogenous gut microbiome in COVID-19 cases compared to healthy controls using systematic, high-resolution multi-omic data, including metagenomics and metatranscriptomics with a particular focus on virulence factors and antimicrobial resistance genes. We find that mild, i.e. asymptomatic-to-moderate, COVID-19 does not alter the overall composition of the gut microbiome, unlike the drastic microbiota changes reported previously in severe cases. Importantly, we find that a mild progression of COVID-19 affects the *infective competence* of gut microbiota, wherein taxa encode and express genes facilitating their survival and/or growth. We find specific families such as Acidaminococcaceae and Erysipelatoclostridiaceae to be encoding for and expressing virulence factors and ARGs, significantly more in individuals with COVID-19. Collectively our data also demonstrates a significantly higher *infective competence* of the endogenous microbiome, suggesting that infection with SARS-CoV-2 may mediate co-infections in the longer term.

## Methods

### Ethics

The study was performed in accordance with the Good Clinical Practice and the Declaration of Helsinki. All participants in this study provided their written informed consent. The study was approved by the National Research Ethics Committee of Luxembourg (study number 202006/03). All biological samples and data were collected in the frame of Predi-COVID [33] and CON-VINCE studies. The studies were approved by the National Research Ethics Committee of Luxembourg (Predi-COVID study number 202003/07; CON-VINCE study number 202004/01).

### Cohort description and patient involvement

Between May and October 2020, stool samples were collected from 61 participants residing in Luxembourg with COVID-19 confirmed by positive SARS-CoV-2 RT-qPCR (Supp. Table 1) within the framework of the Predi-COVID study [33]. In order to be eligible to participate in the study, an individual must have met all the following criteria: (1) signed informed consent form; (2) individuals ≥ 18 years old with confirmed SARS-CoV-2 infection as determined by PCR, performed by one of the certified laboratories in Luxembourg; and (3) hospitalised or at home. In addition to the criteria specific to the Predi-COVID study, samples were excluded if antibiotic treatment was reported. From the individuals, relevant clinical data was collected using a modified version of the International Severe Acute Respiratory and Emerging Infection Consortium (ISARIC) case report form. The participants to be included in the study were classified using an adapted version of the National Institute of Healthy symptom severity scheme [34]. Subsequently, only asymptomatic to moderate symptoms were reported. Along with the samples from the COVID-19 confirmed participants, stool samples were collected from a group of 57 individuals who tested negative for SARS-CoV-2 by RT-qPCR, who were participants of the CON-VINCE study, a population-based cohort study which recruited a representative sample of the Luxembourg population, to serve as age-matched controls. Participation in the control group was excluded if matching any of the following criteria: (1) infection of SARS-CoV-2 prior to the study; (2) presence of fever and respiratory distress/cough not attributable to other known chronic disease; (3) usage of antibiotics up to three months prior to enrolment or first SARS-CoV-2 infection. The study design is presented in Fig. 1 while additional metadata are included in Supp. Table 1. Patients were not involved in setting the research questions or the outcome measures of this study.

**Figure 1.**
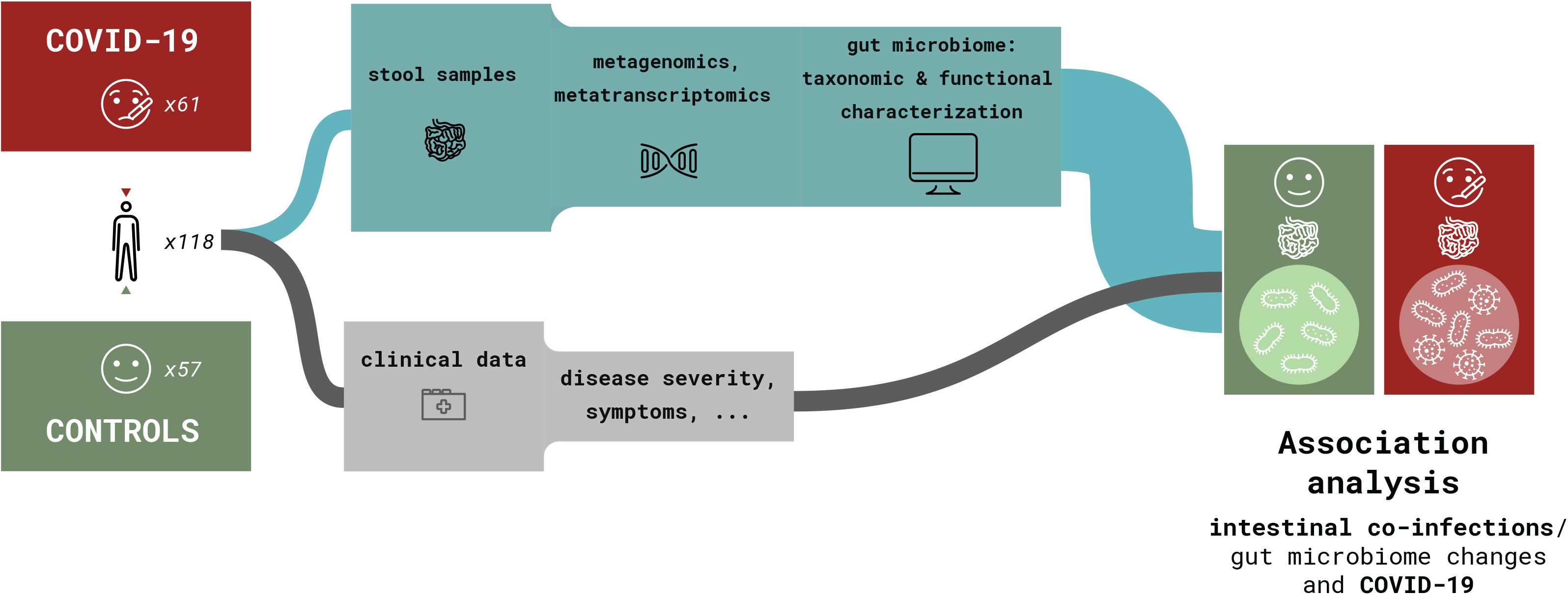
Sample collection and study design. Schematic of the project study design, including cohort composition, and data analyses.

### Sample collection, data processing and prediction of *infective competence*

Stool samples were collected at home by individuals using DNA/RNA Shield Fecal Collection Tubes (Zymo Research), and samples and data were collected at the Integrated BioBank of Luxembourg. DNA and RNA were extracted from all collected stool samples and sequenced for metagenomic and metatranscriptomic analysis. Sequencing was performed at LCSB sequencing platform (RRID: SCR_021931) on NextSeq500 instrument using 2×150 bp read length. The data was analysed using the Integrated Meta-omic Pipeline (IMP; v3 - #b6f9da0e for preprocessing and #c04edbe for downstream analyses) [35]. MetaPhlAn3.1 (v3.1.0, database “mpa_v31_CHOCOPhlAn_201901”) was used for taxonomic profiling [36], while virulence factors and ARGs were identified using PathoFact (v1.0; #6fa64961). Detailed description of the DNA/RNA extractions, sequencing methodology, the analyses approaches, tools and their respective parameters are provided in the Supplementary Material.

### Code and data availability

All code used for metagenomic and metatranscriptomic data analysis can be found in the following repositories: https://gitlab.lcsb.uni.lu/ESB/co-infectomics and https://gitlab.lcsb.uni.lu/laura.denies/co-infectomics_r_analysis. Reads filtered against the human genome (hg38), were submitted to the Sequence Read Archive hosted under accession PRJNA890008. Metadata is included in Supp. Table 1 and all other data can be found on Zenodo (https://doi.org/10.5281/zenodo.7192682).

## Results

### Taxonomic and functional profiles indicate minimal changes in COVID-19

COVID-19 studies have reported an altered gut microbiota composition of hospitalised and critical COVID-19 patients. However, limited attention has been paid to milder forms of COVID-19. Thus, we assessed whether gut microbiota composition was altered in COVID-19 individuals compared to healthy controls. Overall, the gut microbiome compositions, based on the alpha- and beta-diversity metrics, of 61 COVID-19 and 57 individuals from the control group were similar (Fig. 1 and Supp. Fig. 1), with an increased abundance of species belonging to the Lachnospiraceae, Ruminococcaceae, Bacteroidaceae and Bifidobacteriaceae families in COVID-19 (Fig. 2a). We found specific taxonomic differences within the metagenomes, such as an increase in the abundance of AM10 47 (Firmicutes phylum), *Prevotella sp.* CAG 520, *Prevotella stercorea* and *Roseburia sp.* CAG 471 in the COVID-19 group (Fig. 2b), along with a decrease in CAG 145 (Firmicutes phylum), *Roseburia faecis* and *Turicibacter sanguinis* (Fig. 2c). Despite these taxonomic differences, we did not observe any significant changes in the overall functional profile of the microbiome between the COVID-19 and control groups.

**Figure 2.**
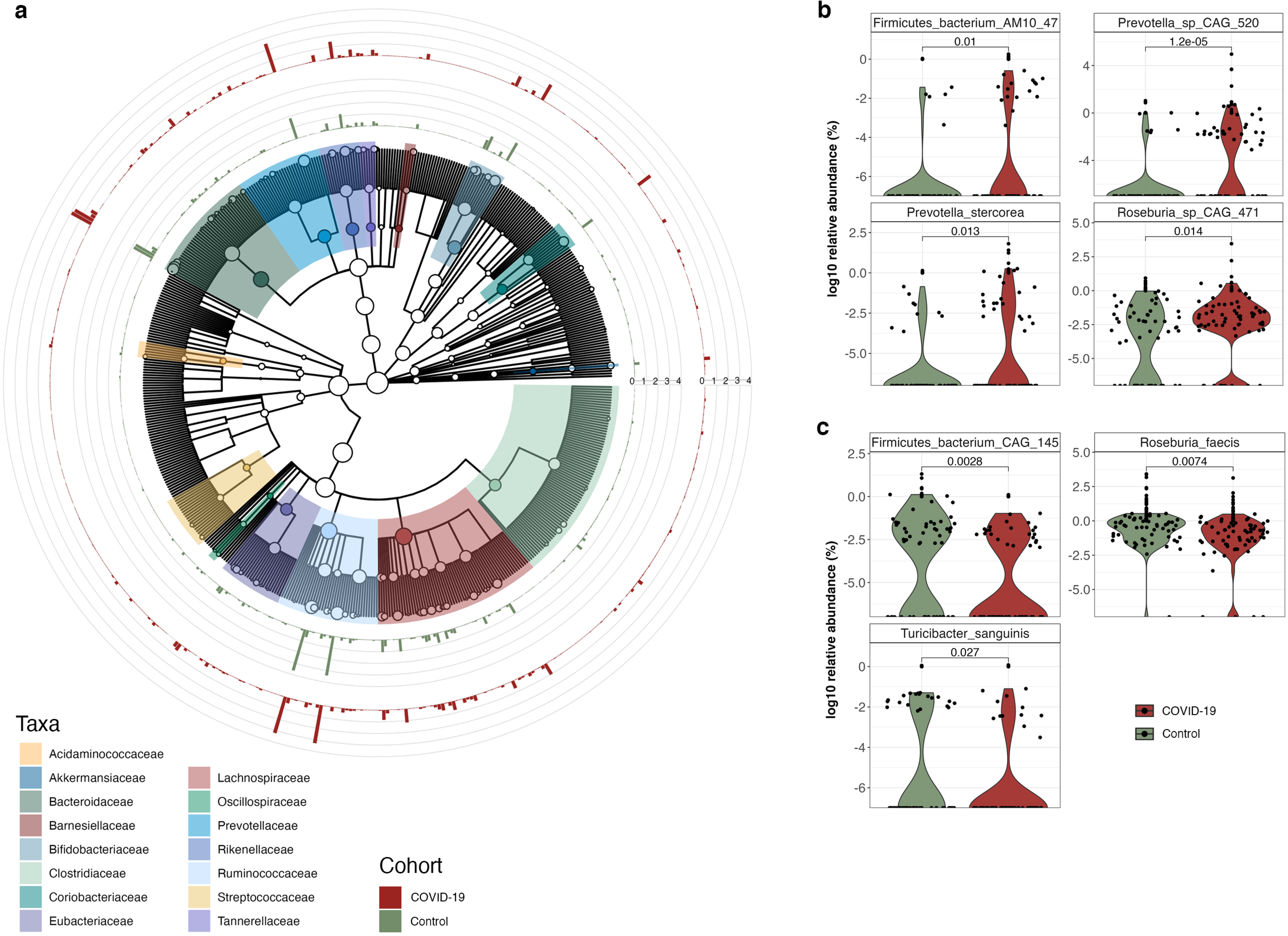
Composition of the microbial community. **a**) Cladogram representing the microbial community profiles in COVID-19 patients (red) and control group (green). The outer rings represent the relative abundance (%) of the microbial community. **b**) Relative abundance of bacterial species significantly enriched in COVID-19 patients compared to the control group [*adj.p* < 0.05; Wilcoxon rank-sum test]. **c**) Relative abundance of bacterial species significantly decreased in COVID-19 patients compared to the control group [*adj.p* < 0.05; Wilcoxon rank-sum test].

In light of reports, indicating the potential co-infections with viruses along with SARS-CoV-2, we also assessed the virome within the COVID-19 patient and the control groups. We did not observe large differences between the groups. However, we found that genes associated with a specific betaherpesvirus and rotavirus were enriched (*adj. p* < 0.05; One-Way ANOVA) in the COVID-19 group (Supp. Table 2).

### SARS-CoV-2 is associated with increased abundance and expression of virulence factors

SARS-CoV-2 infections have been suggested to predispose patients to co-infections or secondary infections of the respiratory and gastrointestinal tracts. Virulence factors in particular enable (pathogenic) microorganisms to colonise host niches and establish infections. We used PathoFact [37] to assess the prevalence of virulence factors in the co-assembled metagenomic and metatranscripomic data. PathoFact was designed to contextualise the genomic data and classify virulence factors and ARGs, allowing to assess the *infective competence* of taxa. To obtain a comprehensive overview of actual gene expression, we complemented metagenomic analyses with metatranscriptomic information conferring information regarding the transcription levels of identified virulence factors. Based on the metagenomic data, we found a significant increase (*adj. p* < 0.05; Wilcoxon rank-sum test) in the overall abundance of virulence factors in the COVID-19 group compared to the control group (Fig. 3a). The metatranscriptomic information further confirmed that these virulence factors demonstrated significantly increased expression levels (*adj. p* < 0.05; Wilcoxon rank-sum test) in the COVID-19 group compared to the control group (Fig. 3b).

**Figure 3.**
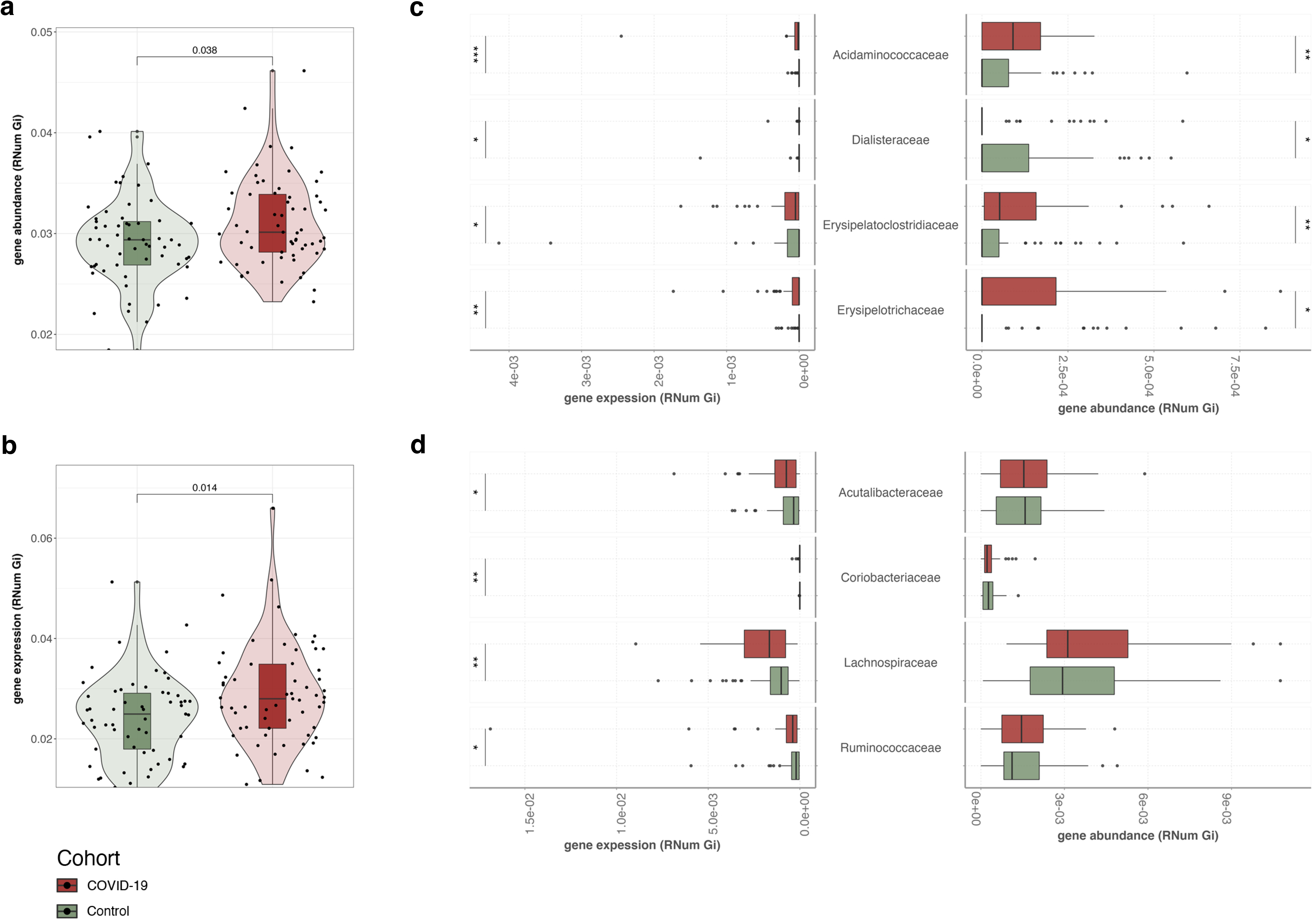
Abundance of virulence factors in the microbial community. **a**) Overall abundance (metagenome) of virulence factors encoded by the microbiome of COVID-19 patients and control group. The significance of the differential abundance is indicated with the adjusted *p*-value [*adj.p* < 0.05; Wilcoxon rank-sum test]. **b**) Overall expression levels (metatranscriptomics) of virulence factors encoded by the microbiome in COVID-19 patients and the control group [*adj.p* < 0.05; Wilcoxon rank-sum test]. **c**) Abundance and expression levels of MAG families where a significant increase in encoded and expressed virulence factors was observed in COVID-19 patients [*adj.p* < 0.05; Wilcoxon rank-sum test, * < 0.05, ** < 0.01, *** < 0.001]. **d**) Abundance and expression levels of virulence factors in MAGs depicting taxonomic families only demonstrating an increased expression of virulence factors, with no significant difference observed at a metagenomic level [*adj.p* < 0.05; Wilcoxon rank-sum test, * < 0.05, ** < 0.01, *** < 0.001].

To link the prevalence and expression of the identified virulence factors to the taxa within the microbial community, we reconstructed metagenome-assembled genomes (MAGs) and further leveraged the iterative workflow of the integrated meta-omic pipeline (IMP) [35]. Overall we found a significant increase in encoded and expressed virulence factors between the COVID-19 and control groups (*adj. p* < 0.05; Wilcoxon rank-sum test). Our analyses further linked families such as Acidaminococcaceae, Erysipelatoclostridiaceae and Erysipelotrichaceae with increased expression of virulence factors in the COVID-19 group (Fig. 3c). Interestingly, the control group exhibited higher gene abundances and expression of virulence factors only in the Dialisteraceae family. Furthermore, we found that some families (Acutalibacteraceae, Coriobacteriaceae, Lachnospiraceae and Ruminococcaceae) demonstrated an increased expression of virulence factors in the COVID-19 group (Fig. 3d; *adj. p* < 0.05; Wilcoxon rank-sum test), although their respective gene abundances were not different from those found in the control group.

### Expression of antimicrobial resistance increases together with virulence factors

While co-infections or secondary infections in COVID-19 may exacerbate the disease, the presence of ARGs may limit treatment options. Since the overall abundance and expression of virulence factors was increased in COVID-19 individuals, we assessed the antimicrobial resistance profile of the microbial community in the COVID-19 and control groups. Specifically, using PathoFact, we characterised the prevalence and expression of ARGs (22 categories). While we did not find any significant differences in the overall gene abundances of all ARGs contributing to the resistome, we observed a significant increase (*adj.p* < 0.05; Wilcoxon rank-sum test) in ARG expression levels between COVID-19 and the control groups (Fig. 4a). Importantly, when investigating individual AMR categories, we found that beta-lactam and peptide resistance were both significantly higher in terms of gene abundance and also more highly expressed within the COVID-19 group (Fig. 4b; *adj.p* < 0.05; Wilcoxon rank-sum test). In addition, we observed that the expression of multidrug resistance was enriched (*adj.p* < 0.05; Wilcoxon rank-sum test) in the COVID-19 group, while macrolides, lincosamides and streptogramins (MLS) resistance also exhibited a higher gene abundance in the same group (*adj.p* < 0.05; Wilcoxon rank-sum test).

**Figure 4.**
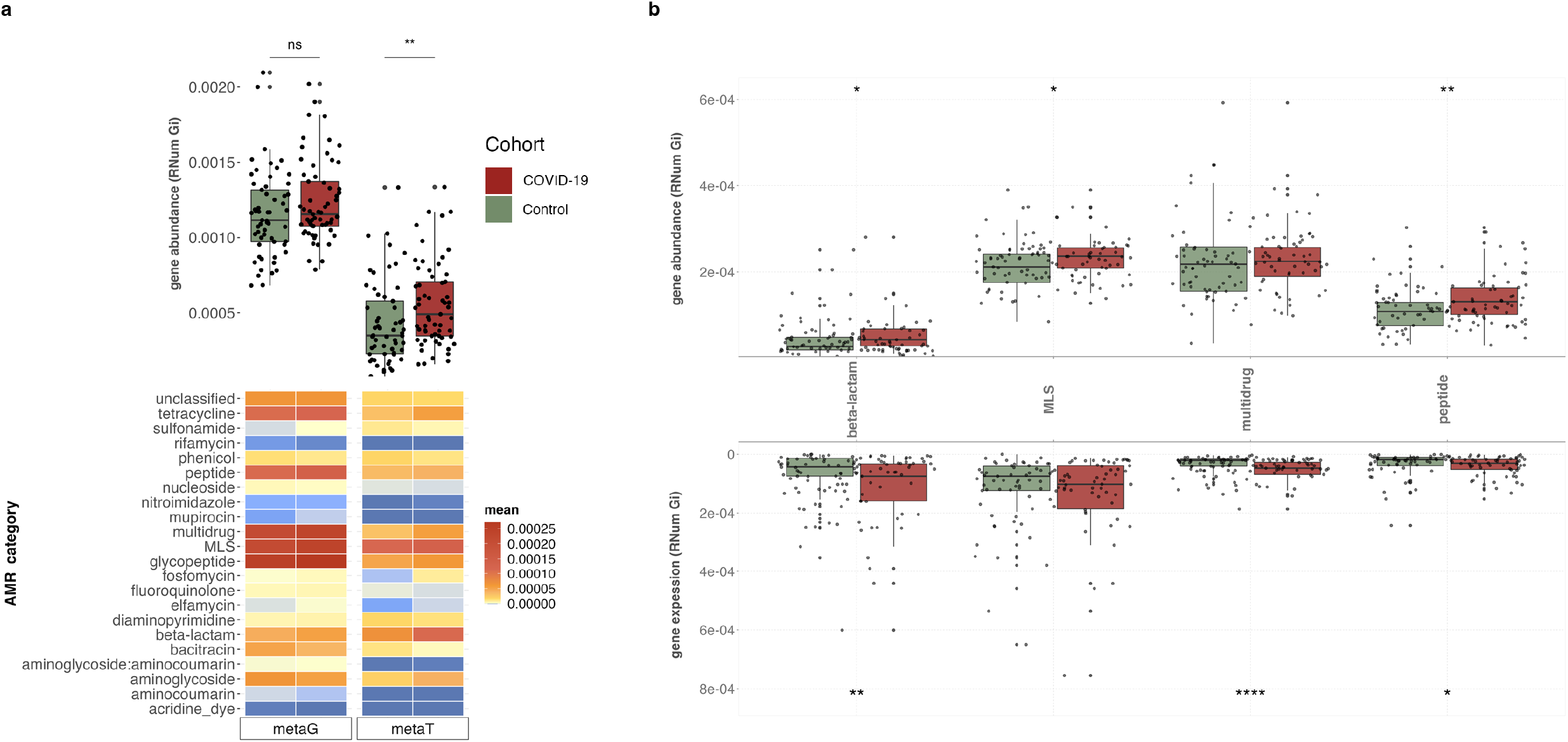
Abundance levels of antimicrobial resistance genes. **a**) Overall ARG abundance and expression levels for COVID-19 and control groups (boxplot), coupled with a breakdown of the respective abundance and expression levels to individual AMR categories [*adj.p* < 0.05; Wilcoxon rank-sum test, * < 0.05, ** < 0.01, *** < 0.001]. **b**) ARG abundance (top) and expression levels (bottom) of individual AMR categories significantly increased in COVID-19 patients compared to the control group [*adj.p* < 0.05; Wilcoxon rank-sum test, * < 0.05, ** < 0.01, *** < 0.001].

As described above, we leveraged the MAGs to correlate the differentially abundant and expressed ARGs to the microbial community. In line with our observations with the virulence factors, we found a significant increase (*adj.p* < 0.05; Wilcoxon rank-sum test) in ARGs encoded and expressed by the Acidaminococcaceae and Erysipelatoclostridiaceae in the COVID-19 group (Fig. 5a-b). Furthermore, an additional family, i.e. Tannerellaceae was also associated with increased abundance and expression of ARGs in the COVID-19 group (Fig. 5a-b). Specifically, in relation to the above reported AMR categories we identified a significant increase in multidrug resistance encoded and expressed by all three of these taxonomic families. In addition, the Acidaminococcaceae also were found to encode a significant increase in ARGs contributing to peptide resistance. Interestingly, we found that several other taxonomic families were also associated with increased ARG expression in the COVID-19 group (*adj.p* < 0.05; Wilcoxon rank-sum test), although their gene abundances did not demonstrate any significant differences (Fig. 5b). These included Barnesiellaceae, Coriobacteriaceae, Lachnospiraceae and Rikenellaceae.

**Figure 5.**
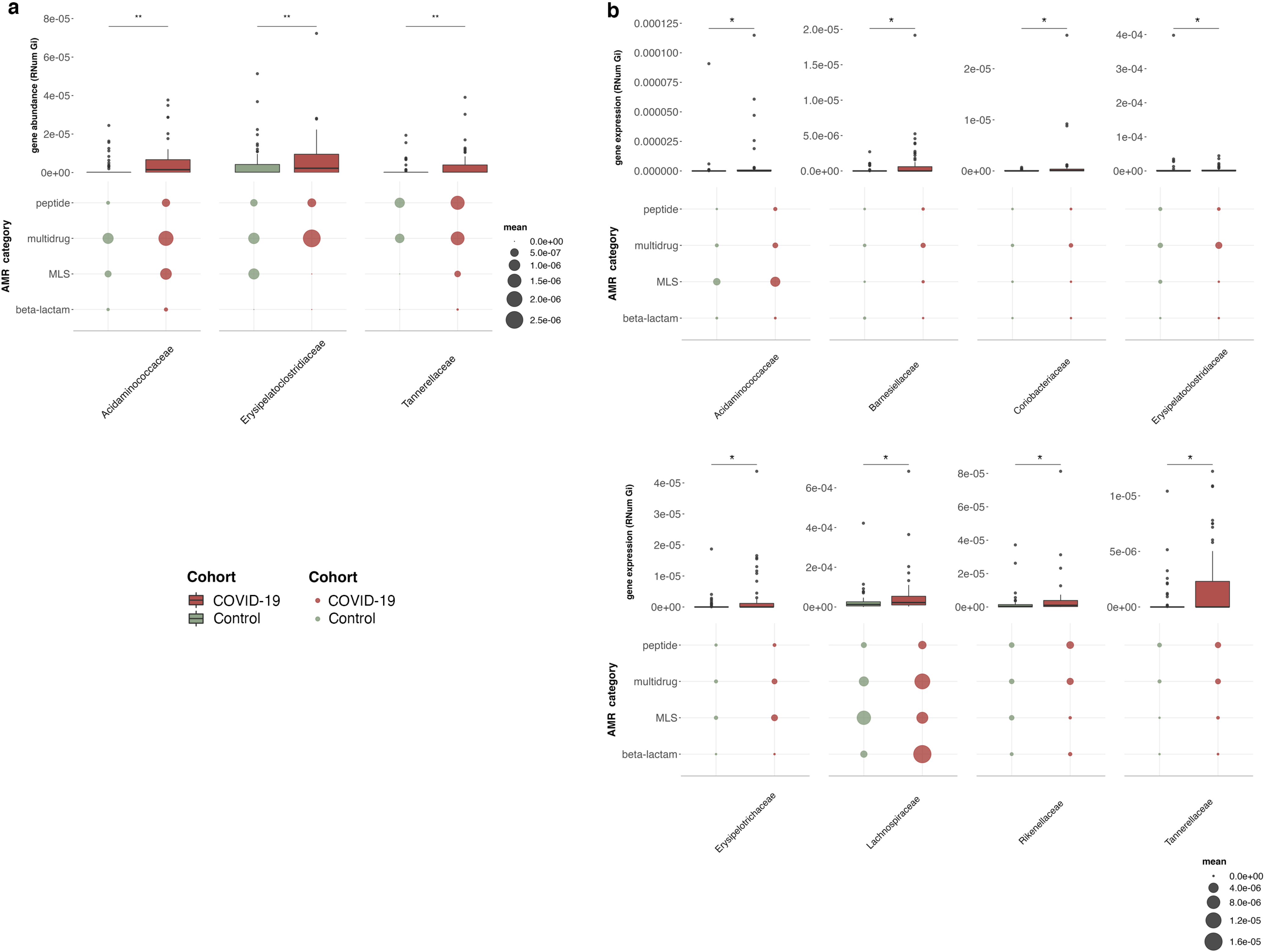
Association of AMR with the microbial community. Abundance (**a**) and expression (**b**) levels of ARGs and corresponding to AMR categories linked to MAGs. On top (boxplot) depicting the overall ARG abundance, below the average abundance of selected AMR categories per taxonomic family. The plot depicts taxonomic families in which overall a significant increase in abundance or expression of ARGs was observed [*adj.p* < 0.05; Wilcoxon rank-sum test, * < 0.05, ** < 0.01, *** < 0.001].

### Infective competence of the gut microbiome

Our analyses collectively indicated that both virulence factors and ARGs were enriched in abundance and expression in the COVID-19 group. Specifically, we found that the abundances of ARGs were correlated with those of the virulence factors (Fig. 6a, R=0.52 and *p* < 0.01; Spearman’s correlation). Complementing this observation, we found that the expression profiles of ARGs and virulence factors also correlated with each other (Fig. 6b, R=0.46 and *p* < 0.01; Spearman’s correlation) suggesting a higher propensity for infectious capacity. To further characterise the *infective competence* of the various taxa within the gut microbiome, we estimated the log2 fold-change of the abundance and expression of virulence factors and ARGs across taxonomic families found in the COVID-19 group and the control group. We found that ~62% (21/34) of the families had a higher *infective competence* and were enriched in abundance and expression within the COVID-19 group, whereas only ~9% (3/34) of the families showed increased *infective competence* in the control group (Fig. 6c). In particular, these analyses highlighted the Acidaminococcaceae and Erysipelatoclostridiaceae families, in line with our earlier observations, suggesting a higher *infective competence*, where the abundances and expression levels of the virulence factors and ARGs were significantly higher in COVID-19 compared to the control group (*p* < 0.05; Two-way ANOVA). In the control group, Dialisteraceae, which was also observed earlier, showed increased *infective competence* (Fig. 6c). Collectively, our data suggests that the *infective competence* of taxa found in the COVID-19 group is increased compared to controls.

**Figure 6.**
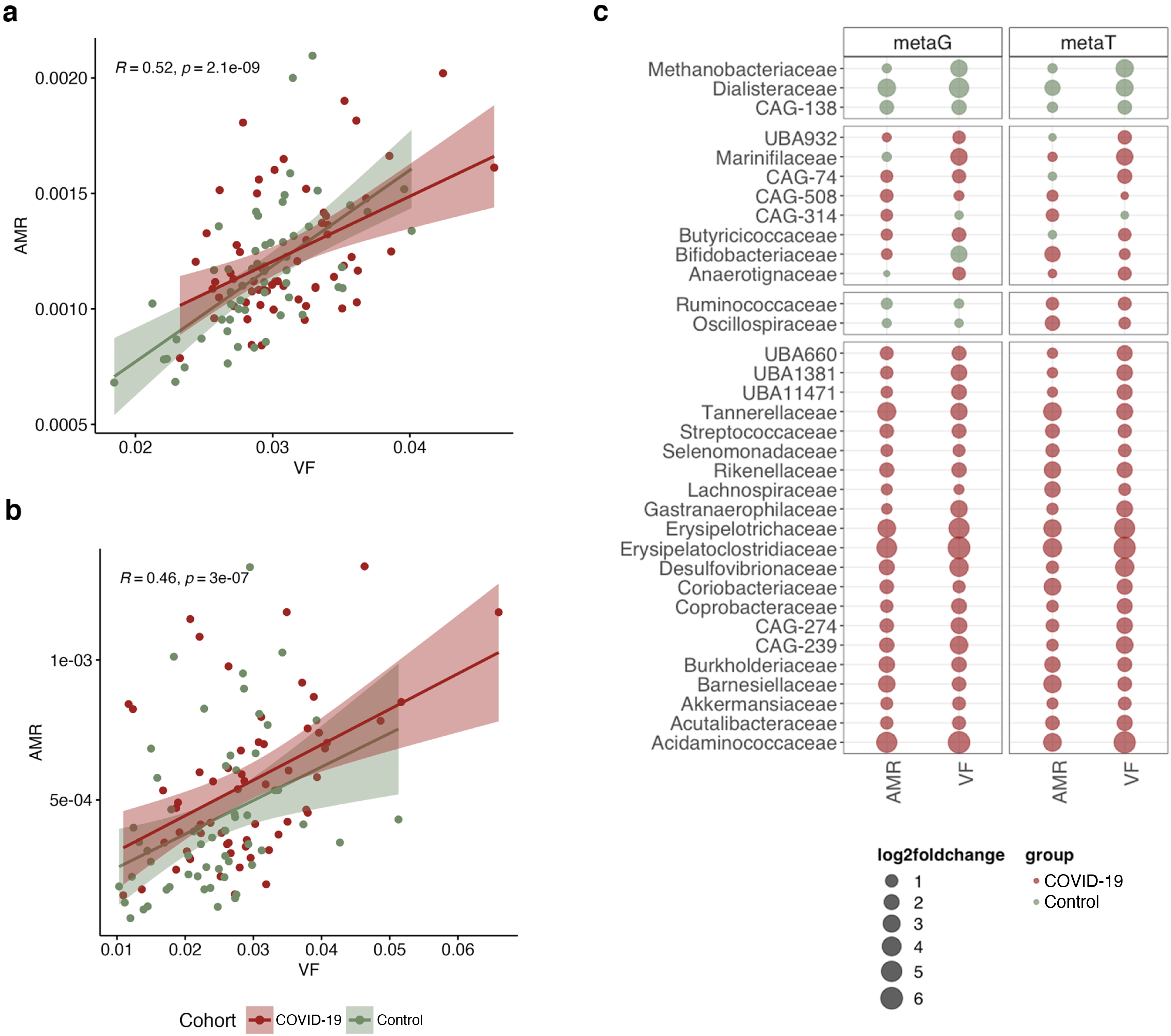
Assessing the *infective competence* of the gut microbiome. **a**) Correlation of gene abundances of AMR and virulence factors [R=0.52 and *p* < 0.01; Spearman’s correlation] in COVID-19 patients (red) and the control group (green). **b**) Correlation of AMR and virulence factors gene expression levels [R=0.46 and *p* < 0.01; Spearman’s correlation] in COVID-19 patients (red) and negative controls (green). **c**) Bubble plot depicting the *infective competence* via the log2 fold change of AMR and virulence factors between COVID-19 patients (red) and control group (green).

## Discussion

COVID-19 has become a common condition for which the manifold effects however remain a challenge [38]. Since the onset of the pandemic, the presentation of gastrointestinal symptoms has indicated the involvement of the gastrointestinal tract in COVID-19 [7]. As we uncover and understand the potential effects of COVID-19 in humans, it is important to also elucidate the concomitant consequences of the disease on the gut microbiome. To this end, several studies have focused on the drastic shifts in the microbiome of COVID-19 patients with severe symptoms. These include changes in diversity [39] including stark enrichments and/or loss of specific taxa [40]. Several Studies have focused on differences in the gut microbiome between patients with severe COVID-19 and controls [7,41,42]. Though these findings are essential, the effect on the larger population, wherein the infection is asymptomatic-to-moderate, is not readily represented. To address this particular gap in knowledge, we focused on the effect of COVID-19 in cases with asymptomatic-to-moderate symptoms in comparison to controls. Interestingly, we found that the diversity and overall shifts in community composition, highlighted in previous reports between severe and control patients [39], did not manifest themselves when comparing asymptomatic-to-moderate cases to the control group of individuals. However, this was associated with an increased abundances of specific taxa such as *Prevotella* spp., AM10 and CAG145 (Firmicutes phylum), *Roseburia* spp. and a *Turicibacter* spp. in the COVID-19 group. This is in contrast to existing reports [40,43], suggesting the loss of beneficial taxa such as *Faecalibacterium*, *Bifidobacterium* and *Roseburia* in the context of COVID-19. Since our study did not include patients with severe COVID-19 or those that were hospitalised, it is likely that the lower disease severity does not lead to significant changes in the abundance of beneficial commensals. Along similar lines, major differences in the virome profile of the COVID-19 group in our study were not observed when compared to the control group. Nevertheless, and importantly, we found that genes associated with rotavirus C were increasingly expressed in the COVID-19 group, despite no differences in overall abundance of this virus between the patient groups. Rotavirus is a known enteric pathogen causing gastroenteritis in the paediatric population, however, their capacity to cause infections in adults is underappreciated and poorly characterised due to only mild symptoms including nausea, headaches and diarrhoea [44]. Importantly, at the time of writing only one report by Wang et al. [45], indicated the possibility of increased rotavirus A-mediated enteric infections in COVID-19 patients. These findings are intriguing given the propensity for COVID-19 patients to suffer from enteric symptoms [3], including nausea [46] and diarrhoea [47]. Whether the rotavirus, especially in adults, is associated with COVID-19 gastrointestinal symptoms, or the enteric effects exacerbate the expression of rotavirus C-associated genes is still unknown and will have to be investigated in dedicated follow-up studies.

In line with the above observations, early in the pandemic, the role of COVID-19 in enhancing co-infections was documented extensively [19,48,49]. This is not limited to mucormycosis [50] which was amplified in certain parts of the world, but also bacterial and viral co-infections that were reported in severe COVID-19 patients [45]. Despite these observations and case studies, the effect of COVID-19 on the *infective competence* of the existing and endogenous microbiota has never been characterised. Our findings, therefore, bridge an important and broad chasm in knowledge, suggesting that COVID-19-mediated shifts may lead to higher microbiome-linked burden with potentially manifold effects. Importantly, we not only found an increased abundance in virulence factors in the COVID-19 group, but also a concomitant increase in expression of genes associated with virulence. Such infection-linked shifts have not been reported beforehand and may be used to monitor and understand potential future infections, in the context of COVID-19-mediated effects. Simultaneously, we found that these virulence factors were associated with taxa from families such as Acidaminococcaceae, Erysipelatoclostridiaceae and Erysipelotrichaceae. Though Acidaminococcus was recently reported to be associated with a disease-related group in a large-scale meta-analysis [51], the exact role of Acidaminococcaceae in virulence is undocumented. Members of the Erysipelatoclostridiaceae family are typically seen as typical members of the microbiome, however in specific cases species such as *Erysipelatoclostrium ramosum* have been associated with systemic infection and systemic inflammatory response syndrome [52], while Erysipelotrichaceae have been positively correlated with colorectal cancer [53]. Our observations, especially the increased expression of virulence factor genes associated with these taxa, may pave the way in future explorations to serve as indicators of diseases. Importantly, it is still unclear whether the enriched *infective competence* is a COVID-19-specific hallmark or one found in all infections. For example, it has previously been hypothesised that selection of pathobionts result from inflammatory responses and/or a dysregulation of the tolerant immune system [54,55]. Future studies will need to address the extent to which various underlying factors such as a dysbiotic microbiome and an impaired immune system affect the *infective competence* of the gut microbiome.

Another important aspect of COVID-19, in particular early on in the pandemic, was the overuse and misuse of antibiotics for treating SARS-CoV-2 [56,57] which was also associated with the potential increase of AMR [23,58]. Recent studies have reported on the higher incidence of AMR [58] and increased ARGs in COVID-19 patients [59,60]. However, these reports either refer to patients who were administered antibiotics [59] or include a meta-analyses observing datasets which were generated pre- and post-pandemic, specifically associated with travel [61], or limit characterisation of antibiotic-mediated differences at a broad and low-resolution [62]. In contrast to these studies, antibiotic usage was a clear exclusion criterion in our study where individuals included were not administered any antibiotics three months prior to sampling. To our knowledge, our findings are the first report to systematically analyse the resistome of COVID-19 and control individuals and importantly to demonstrate that several of these ARGs are indeed expressed significantly higher in the COVID-19 group compared to the control group, regardless of antibiotic treatment. We observe that resistance genes include MLS, multidrug and peptides, resistance classes where treatments of resistant bacteria are known to be inherently challenging with conventional antibiotics [63,64]. Strikingly, we found that the increased ARG expression in the COVID-19 group was further associated with the same taxa encoding and expressing virulence factors. This suggests that combinatorial effects of virulence factors and ARGs may exacerbate the *infective competence* of these taxa. This is further supported by our analysis identifying that taxa from the Acidaminococcaceae and Erysipelatoclostridiaceae families demonstrated a predicted higher *infective competence* in the COVID-19 group.

Our findings suggest that it is imperative to elucidate all the implications of SARS-CoV-2 infection, especially its effect on the gut microbiome community and functions. Although other studies have involved the severe cases of COVID-19 [7,65,66], none of these studies include both metagenomic and metatranscriptomic sequencing data. We found that the virulence factors and antimicrobial resistance genes were indeed expressed in higher levels in the COVID-19 group compared to the controls. These key findings would have not been possible by only focussing on metagenomic data. Our collective findings, indicating the enriched abundance and expression of both virulence factors and ARGs, suggest that COVID-19 may yet have unknown effects that may come to light in the longer term including the shaping of the microbiome across the population. Moreover, we find that none of the commonly reported pathogens *(Salmonella*, *Shigella*, *Klebsiella* etc.) are enriched in the COVID-19 group in our study. In contrast, we find changes in *Prevotella* spp, AM10 and CAG145 (Firmicutes phylum), *Roseburia* spp and a *Turicibacter* spp. Therefore, it will be critically important to evaluate and further validate the effects of COVID-19 on the gut microbiome also in relation to infections by other viral and other pathogens. In particular, it remains unclear at this time, whether infections with other viruses, known to cause respiratory and gastrointestinal distress, e.g. Adenoviruses, respiratory syncytial virus (RSV), influenza viruses, norovirus, would lead to similar community and functional changes within the gut microbiome. Overall, it must be reiterated that pandemic preparedness coupled to the monitoring of virulence factors in tandem with antibiotic stewardship may be essential components for future strategies to mitigate the longer-term effects of COVID-19 and possibly other viral infections.

## Acknowledgements

We acknowledge the active involvement of all the participants of the Predi-COVID and CON-VINCE cohorts. We are also thankful for the support of the cohort recruitment and processing teams. We also acknowledge Prof. Laetitia Huiart for her efforts in initiating the Predi-COVID cohort and Aurélie Sausy, Sophie Mériaux, and Jean-Yves Servais for their technical help. The Predi-COVID study is supported by the Luxembourg National Research Fund (FNR: 14716273), the Fondation André Losch, and the European Regional Development Fund (FEDER, no. 2018-04-026-21). F.Q.H. was partially supported by FNR AFR-RIKEN bilateral program (TregBAR, 11228353, F.Q.H. and M.O.) and PRIDE programme (11012546/NEXTIMMUNE and 10907093/CRITICS). The CON-VINCE study is supported by the Fonds National de la Recherche (FNR: 14716281/CON-VINCE/Kruger) and the André Losch Foundation. P.W. acknowledges funding from the European Research Council under the European Union’s Horizon 2020 research and innovation program (no. 863664). LdN, PM and PW are supported by the Luxembourg National Research Fund (FNR; PRIDE17/11823097) awarded to PW. We would also like to acknowledge the following funding instruments: FNR: COVID-19/2020-1/14719552/FunBiome. The work was further supported by the Luxembourg Government through the CoVaLux programme. This research has also been supported by the Swiss National Science Foundation (CRSII5_180241) supporting SBB. We would like to thank Susana Martinez Arbas for the initial support during data preprocessing. The authors are also grateful to the Luxembourg Ministry of Higher Education and Research and the Ministry of Health for funding the intervention. The computational analyses were performed at the HPC facilities at the University of Luxembourg (https://hpc.uni.lu) [67].

## Authors’ contributions

The authors are solely responsible for all aspects of this article including the research, the interpretation, and the writing thereof. PW, LD, CMG, SBB and CCL conceived the study, while LdN, VG and SBB performed data analysis, data interpretation and in conjunction with MD wrote the manuscript. CS, FH, and PM contributed to critical experiments and analyses. RH, JH and PCL were instrumental in sequencing the samples. GF, MO, JVF, TM, ES, MPOS, SG, VS were responsible for the data and sample collection of the Predi-COVID or CON-VINCE cohorts. All authors were involved in writing and approval of the manuscript.

## The CON-VINCE Consortium^†^

Geeta Acharya, Luxembourg Institute of Health, Strassen, Luxembourg; Gloria Aguayo, Luxembourg Institute of Health, Strassen, Luxembourg; Wim Ammerlaan, Luxembourg Institute of Health, Strassen, Luxembourg; Ariane Assele-Kama, Luxembourg Institute of Health, Strassen, Luxembourg; Christelle Bahlawane, Luxembourg Institute of Health, Strassen, Luxembourg; Katy Beaumont, Luxembourg Institute of Health, Strassen, Luxembourg; Nadia Beaupain, Luxembourg Institute of Health, Strassen, Luxembourg; Lucrèce Beckers, Luxembourg Institute of Health, Strassen, Luxembourg; Camille Bellora, Luxembourg Institute of Health, Strassen, Luxembourg; Fay Betsou, Luxembourg Institute of Health, Strassen, Luxembourg; Sandie Boly, Luxembourg Institute of Health, Strassen, Luxembourg; Dirk Brenner, Luxembourg Institute of Health, Strassen, Luxembourg; Eleftheria Charalambous, Luxembourg Institute of Health, Strassen, Luxembourg; Emilie Charpentier, Luxembourg Institute of Health, Strassen, Luxembourg; Manuel Counson, Luxembourg Institute of Health, Strassen, Luxembourg; Brian De Witt, Luxembourg Institute of Health, Strassen, Luxembourg; Olivia Domingues, Luxembourg Institute of Health, Strassen, Luxembourg; Claire Dording, Luxembourg Institute of Health, Strassen, Luxembourg; Bianca Dragomir, Luxembourg Institute of Health, Strassen, Luxembourg; Tessy Fautsch, Luxembourg Institute of Health, Strassen, Luxembourg; Jean-Yves Ferrand, Luxembourg Institute of Health, Strassen, Luxembourg; Ana Festas Lopes, Luxembourg Institute of Health, Strassen, Luxembourg; Joëlle Véronique Fritz, Luxembourg Institute of Health, Strassen, Luxembourg; Manon Gantenbein, Luxembourg Institute of Health, Strassen, Luxembourg; Laura Georges, Luxembourg Institute of Health, Strassen, Luxembourg; Jérôme Graas, Luxembourg Institute of Health, Strassen, Luxembourg; Gael Hamot, Luxembourg Institute of Health, Strassen, Luxembourg; Anne-Marie Hanff, Luxembourg Institute of Health, Strassen, Luxembourg; Maxime Hansen, Luxembourg Institute of Health, Strassen, Luxembourg; Lisa Hefele, Luxembourg Institute of Health, Strassen, Luxembourg; Estelle Henry, Luxembourg Institute of Health, Strassen, Luxembourg; Margaux Henry, Luxembourg Institute of Health, Strassen, Luxembourg; Eve Herkenne, Luxembourg Institute of Health, Strassen, Luxembourg; Christiane Hilger, Luxembourg Institute of Health, Strassen, Luxembourg; Judith Hübschen, Luxembourg Institute of Health, Strassen, Luxembourg; Laetitia Huiart, Luxembourg Institute of Health, Strassen, Luxembourg; Alexander Hundt, Luxembourg Institute of Health, Strassen, Luxembourg; Gilles Iserentant, Luxembourg Institute of Health, Strassen, Luxembourg; Stéphanie Kler, Luxembourg Institute of Health, Strassen, Luxembourg; Rejko Krüger, Luxembourg Institute of Health, Strassen, Luxembourg; Pauline Lambert, Luxembourg Institute of Health, Strassen, Luxembourg; Sabine Lehmann, Luxembourg Institute of Health, Strassen, Luxembourg; Morgane Lemaire, Luxembourg Institute of Health, Strassen, Luxembourg; Andrew Lumley, Luxembourg Institute of Health, Strassen, Luxembourg; Monica Marchese, Luxembourg Institute of Health, Strassen, Luxembourg; Sophie Mériaux, Luxembourg Institute of Health, Strassen, Luxembourg; Maura Minelli, Luxembourg Institute of Health, Strassen, Luxembourg; Alessandra Mousel, Luxembourg Institute of Health, Strassen, Luxembourg; Maeva Munsch, Luxembourg Institute of Health, Strassen, Luxembourg; Mareike Neumann, Luxembourg Institute of Health, Strassen, Luxembourg; Markus Ollert, Luxembourg Institute of Health, Strassen, Luxembourg; Marc Paul O’Sullivan, Luxembourg Institute of Health, Strassen, Luxembourg; Magali Perquin, Luxembourg Institute of Health, Strassen, Luxembourg; Achilleas Pexaras, Luxembourg Institute of Health, Strassen, Luxembourg; Jean-Marc Plesseria, Luxembourg Institute of Health, Strassen, Luxembourg; Lucie Remark, Luxembourg Institute of Health, Strassen, Luxembourg; Estelle Sandt, Luxembourg Institute of Health, Strassen, Luxembourg; Bruno Santos, Luxembourg Institute of Health, Strassen, Luxembourg; Aurélie Sausy, Luxembourg Institute of Health, Strassen, Luxembourg; Margaux Schmitt, Luxembourg Institute of Health, Strassen, Luxembourg; Sneeha Seal, Luxembourg Institute of Health, Strassen, Luxembourg; Jean-Yves Servais, Luxembourg Institute of Health, Strassen, Luxembourg; Florian Simon, Luxembourg Institute of Health, Strassen, Luxembourg; Chantal Snoeck, Luxembourg Institute of Health, Strassen, Luxembourg; Kate Sokolowska, Luxembourg Institute of Health, Strassen, Luxembourg; Hermann Thien, Luxembourg Institute of Health, Strassen, Luxembourg; Johanna Trouet, Luxembourg Institute of Health, Strassen, Luxembourg; Jonathan Turner, Luxembourg Institute of Health, Strassen, Luxembourg; Michel Vaillant, Luxembourg Institute of Health, Strassen, Luxembourg; Daniela Valoura Esteves, Luxembourg Institute of Health, Strassen, Luxembourg; Charlène Verschueren, Luxembourg Institute of Health, Strassen, Luxembourg; Tania Zamboni, Luxembourg Institute of Health, Strassen, Luxembourg; Pinar Alper, Luxembourg Centre for Systems Biomedicine, University of Luxembourg, Esch-Belval, Luxembourg; Piotr Gawron, Luxembourg Centre for Systems Biomedicine, University of Luxembourg, Esch-Belval, Luxembourg; Soumyabrata Ghosh, Luxembourg Centre for Systems Biomedicine, University of Luxembourg, Esch-Belval, Luxembourg; Enrico Glaab, Luxembourg Centre for Systems Biomedicine, University of Luxembourg, Esch-Belval, Luxembourg; Clarissa Gomes, Luxembourg Centre for Systems Biomedicine, University of Luxembourg, Esch-Belval, Luxembourg; Borja Gomez Ramos, Luxembourg Centre for Systems Biomedicine, University of Luxembourg, Esch-Belval, Luxembourg; Vyron Gorgogietas, Luxembourg Centre for Systems Biomedicine, University of Luxembourg, Esch-Belval, Luxembourg; Valentin Groues, Luxembourg Centre for Systems Biomedicine, University of Luxembourg, Esch-Belval, Luxembourg; Wei Gu, Luxembourg Centre for Systems Biomedicine, University of Luxembourg, Esch-Belval, Luxembourg; Laurent Heirendt, Luxembourg Centre for Systems Biomedicine, University of Luxembourg, Esch-Belval, Luxembourg; Ahmed Hemedan, Luxembourg Centre for Systems Biomedicine, University of Luxembourg, Esch-Belval, Luxembourg; Sascha Herzinger, Luxembourg Centre for Systems Biomedicine, University of Luxembourg, Esch-Belval, Luxembourg; Anne Kaysen, Luxembourg Centre for Systems Biomedicine, University of Luxembourg, Esch-Belval, Luxembourg; Jacek Jaroslaw Lebioda, Luxembourg Centre for Systems Biomedicine, University of Luxembourg, Esch-Belval, Luxembourg; Tainà Marques, Luxembourg Centre for Systems Biomedicine, University of Luxembourg, Esch-Belval, Luxembourg; François Massart, Luxembourg Centre for Systems Biomedicine, University of Luxembourg, Esch-Belval, Luxembourg; Patrick May, Luxembourg Centre for Systems Biomedicine, University of Luxembourg, Esch-Belval, Luxembourg; Christiane Olesky, Luxembourg Centre for Systems Biomedicine, University of Luxembourg, Esch-Belval, Luxembourg; Venkata P. Satagopam, Luxembourg Centre for Systems Biomedicine, University of Luxembourg, Esch-Belval, Luxembourg; Claire Pauly, Luxembourg Centre for Systems Biomedicine, University of Luxembourg, Esch-Belval, Luxembourg; Laure Pauly, Luxembourg Centre for Systems Biomedicine, University of Luxembourg, Esch-Belval, Luxembourg; Lukas Pavelka, Luxembourg Centre for Systems Biomedicine, University of Luxembourg, Esch-Belval, Luxembourg; Guilherme Ramos Meyers, Luxembourg Centre for Systems Biomedicine, University of Luxembourg, Esch-Belval, Luxembourg; Armin Rauschenberger, Luxembourg Centre for Systems Biomedicine, University of Luxembourg, Esch-Belval, Luxembourg; Basile Rommes, Luxembourg Centre for Systems Biomedicine, University of Luxembourg, Esch-Belval, Luxembourg; Kirsten Rump, Luxembourg Centre for Systems Biomedicine, University of Luxembourg, Esch-Belval, Luxembourg; Reinhard Schneider, Luxembourg Centre for Systems Biomedicine, University of Luxembourg, Esch-Belval, Luxembourg; Valerie Schröder, Luxembourg Centre for Systems Biomedicine, University of Luxembourg, Esch-Belval, Luxembourg; Amna Skrozic, Luxembourg Centre for Systems Biomedicine, University of Luxembourg, Esch-Belval, Luxembourg; Lara Stute, Luxembourg Centre for Systems Biomedicine, University of Luxembourg, Esch-Belval, Luxembourg; Noua Toukourou, Luxembourg Centre for Systems Biomedicine, University of Luxembourg, Esch-Belval, Luxembourg; Christophe Trefois, Luxembourg Centre for Systems Biomedicine, University of Luxembourg, Esch-Belval, Luxembourg; Carlos Vega Moreno, Luxembourg Centre for Systems Biomedicine, University of Luxembourg, Esch-Belval, Luxembourg; Maharshi Vyas, Luxembourg Centre for Systems Biomedicine, University of Luxembourg, Esch-Belval, Luxembourg; Xinhui Wang, Luxembourg Centre for Systems Biomedicine, University of Luxembourg, Esch-Belval, Luxembourg; Anja Leist, University of Luxembourg, Esch-sur-Alzette, Luxembourg; Annika Lutz, University of Luxembourg, Esch-sur-Alzette, Luxembourg; Claus Vögele, University of Luxembourg, Esch-sur-Alzette, Luxembourg; Linda Hansen, Centre Hospitalier de Luxembourg, Luxembourg, Luxembourg; João Manuel Loureiro, Centre Hospitalier de Luxembourg, Luxembourg, Luxembourg; Beatrice Nicolai, Centre Hospitalier de Luxembourg, Luxembourg, Luxembourg; Alexandra Schweicher, Centre Hospitalier de Luxembourg, Luxembourg, Luxembourg; Femke Wauters, Centre Hospitalier de Luxembourg, Luxembourg, Luxembourg; Tamir Abdelrahman, Laboratoire National de Santé, Dudelange, Luxembourg; Estelle Coibion, Laboratoire National de Santé, Dudelange, Luxembourg; Guillaume Fournier, Laboratoire National de Santé, Dudelange, Luxembourg; Marie Leick, Laboratoire National de Santé, Dudelange, Luxembourg; Friedrich Mühlschlegel, Laboratoire National de Santé, Dudelange, Luxembourg; Marie France Pirard, Laboratoire National de Santé, Dudelange, Luxembourg; Nguyen Trung, Laboratoire National de Santé, Dudelange, Luxembourg; Philipp Jägi, Laboratoire Réunis, Junglinster, Luxembourg; Henry-Michel Cauchie, Luxembourg Institute of Science and Technology, Luxembourg, Luxembourg; Delphine Collart, Luxembourg Institute of Science and Technology, Luxembourg, Luxembourg; Leslie Ogorzaly, Luxembourg Institute of Science and Technology, Luxembourg, Luxembourg; Christian Penny, Luxembourg Institute of Science and Technology, Luxembourg, Luxembourg; Cécile Walczak, Luxembourg Institute of Science and Technology, Luxembourg, Luxembourg.

## Supplementary Material

### Sample collection and processing

Stool samples were collected at home by individuals using DNA/RNA Shield Fecal Collection Tubes (Zymo Research), and samples and data were collected at the Integrated BioBank of Luxembourg (IBBL). Around 1 g of stool was sampled, diluted in 9 ml DNA/RNA Shield according to the manufacturer’s suggestions. Subsequently, stool samples in DNA/RNA shield were thawed on ice and aliquoted according to the following schema: 250 μl of sample was used to extract DNA, 250 μl of lysis solution (ZymoBIOMICS DNA Miniprep Kit; Zymo Research) was added and sample was kept frozen at −80°C until DNA extraction was performed. 700 μl was used to extract RNA using ZR BashingBead Lysis Tubes (Zymo Research) and RNeasy Mini Kit (QIAGEN). Extractions were performed right after thawing.

### DNA and RNA extraction

Samples for DNA extraction were extracted using ZymoBIOMICS DNA Miniprep Kit according to the manufacturer’s instructions with the following modifications: samples were inactivated for 7 min at 70°C prior to homogenization by milling for 3 cycles (5 min of cooling on ice between cycles) for 60 s at 6 m/s in a FastPrep-24 5 G (MP Biomedicals). Prior to DNA purification, a Proteinase K incubation step was performed: 5 μl of 20 mg/ml Proteinase K (New England Biolabs GmbH) was added to each sample and incubated for 30 min at 40°C. Extraction was performed following the supplier’s manual and DNA was eluted in 50 μl DNase/RNase-Free Water (prewarmed to 60°C). RNase treatment was performed by addition of 2.4 μl of 20 mg/ml Monarch RNase A (New England Biolabs GmbH) to each sample and incubated for 10 min at 56°C. DNA was purified and concentrated using ZR-96 DNA Clean-Up Kit (Zymo Research) following the supplier’s manual and DNA was eluted in 50 μl DNase/RNase-Free water (prewarmed to 60°C). DNA was quantified using Qubit dsDNA BR assay kit (Invitrogen) and purity determined using Nanodrop 2000C (Thermo Scientific). Samples were frozen at −80°C for further use.

Samples for RNA extraction were inactivated for 7 min at 70°C and 600 μl of cold RLT Buffer (containing 10 μl/ml 2-mercaptoethanol) was added to the samples prior to homogenization by milling for 3 cycles (5 min of cooling on ice between cycles) for 60 s at 6 m/s in a FastPrep-24 5 G (MP Biomedicals). Samples were centrifuged for 3 min at full speed and supernatant was mixed with 1 volume of 70 % Ethanol. Lysates were loaded onto a RNeasy Mini Spin Column and centrifuged at 8,000 x g for 1 min. This last step was repeated until all supernatants had passed through the filter. Columns were washed according to the supplier’s manual and 50 μl RNase-free water was added to the centre of the filter and incubated at room temperature for 1 min. RNA was eluted by centrifugation at 8,000 × g for 1 min. RNA was filled up to 87.5 μl with RNase-free water, 2.5 μl DNase I stock solution and 10 μl Buffer RDD (both RNase-Free DNase Set, QIAGEN) was added, mixed and incubated for 10 min at room temperature. RNA was purified and concentrated using RNA Clean & Concentrator-5 kit (Zymo Research) following the supplier’s manual. RNA was eluted in 15 μl DNase/RNase-Free water. 1 μl of obtained RNA was heat‐denatured for 2 min at 72°C and quality-checked using Agilent RNA 6000 Nano kit (Agilent Technologies). RNA was quantified using Qubit RNA HS assay kit (Invitrogen). Samples were frozen at −80°C for further use.

### Metagenomic and metatranscriptomic sequencing

DNA and RNA were extracted from all collected stool samples and sequenced for metagenomic and metatranscriptomic analysis. 100ng of DNA was used for metagenomic library preparation using Swift 2S turbo Flexible DNA library kit (cat. no. 45096). The genomic DNA was enzymatically fragmented for 10 min and DNA libraries were prepared without PCR amplification. The average insert size of libraries was 600bp. Prepared libraries were quantified using Qubit (DNA HS kit, ThermoFischer) and quality checked with DNA HS kit on Bioanalyzer 2100 (Agilient). Sequencing was performed at LCSB sequencing platform (RRID: SCR_021931) on NextSeq2000 instrument using 2×150 bp read length.

500 ng of RNA was rRNA depleted using Illumina Ribo-Zero Plus rRNA Depletion kit (Illumina, 20037135). rRNA depleted samples were further prepared using TruSeq Stranded mRNA library preparation kit (Illumina, 20020594) which includes the fragmentation and priming steps. The fragmentation time was reduced to 3 min. Prepared libraries were quantified using Qubit (DNA HS kit, ThermoFischer) and quality checked with DNA HS kit on Bioanalyzer 2100 (Agilient). Sequencing was performed at LCSB sequencing platform (RRID: SCR_021931) on NextSeq500 instrument using 2×150 bp read length. In total this resulted in ~6Gbp per sample for the metagenomics and ~21 Gbp per sample for the metatranscriptomics.

### Data processing, including genome reconstruction

The Integrated Meta-omic Pipeline (IMP; v3 - commitID #b6f9da0e for preprocessing and #c04edbe for downstream assemblies) [39] was used for the processing and iterative co-assembly of metagenomic and metatranscriptomic reads. The workflow includes pre-processing, assembly, genome reconstruction, and functional and taxonomic annotation based on public and custom databases in a reproducible manner. For the data preprocessing, raw metagenomic reads were first trimmed to the maximal read length of 150 bases using Cutadapt (v3.4) [40]. The preprocessed metagenomic and raw metatranscriptomic reads were processed using IMP: reads were trimmed using Trimmomatic (v.39) [41], reads mapping to the human genome (hg38 genome) or PhiX genome (gi|9626372|ref|NC_001422.1, Enterobacteria phage phiX174 sensu lato, complete genome) were removed using BWA (v. 0.7.9a) [42], and the metatranscriptomic reads were further filtered using SortMeRNA (v.4.2.0-238-g90cdf6c) [43]. In addition, alpha-diversity estimation was performed from metagenomic reads using Nonpareil (v. 3.4.1) [44] as part of the IMP preprocessing step. Quality control was done on the processed reads by running FastQC (v. 0.11.9) [45] and summarising the reports using MultiQC (v. 1.10.1) [46]. In addition, Kraken2 (v. 2.1.2) [47] was used with a database containing only the human and PhiX genomes (https://ndownloader.figshare.com/files/24658262, from 11.09.2020, provided by Mike Lee) to confirm the successful removal of these contaminants from the processed sequencing data. To detect other viruses and confirm the status of SARS-CoV-2 infection in the processed reads, we used fastv (v. 0.8.1, v0.8.1 for SARS-CoV-2 data, data for other viruses was downloaded on September 11th, 2021) [48]. The tool bbmap (v. 38.90) [49] was used on the preprocessed FASTQ files to extract reads mapping to SARS-CoV-2 reference genomes (same genomes as provided by fastv). Pairwise sample (dis)similarity was estimated using Mash (v. 2.3) [50].

*De novo* co-assembly of the processed metagenomic and metatranscriptomic reads was performed by running Megahit (v2.0) [51] included in IMP, followed by gene calling using an in-house modified Prokka version also allowing for incomplete ORFs [52]. Concurrently, MetaBAT2 [53] and MaxBin2 [54] together with an in-house binning methodology, binny [55], were used to reconstruct metagenome-assembled genomes (MAGs). Subsequently, we obtained a non-redundant set of MAGs using DAS Tool (v1.1.4) [56] with a score threshold of 0.7 for downstream analyses, and those with a minimum completion of 90% and less than 5% contamination as assessed by CheckM (v1.1.3) [57]. Taxonomy was assigned to the MAGs using gtdbtk (v1.7.0) [58]. Finally, MetaQUAST (v. 5.0.2) [59] was run on the created contig FASTA files to compute assembly statistics such as the number and maximal length of contigs, total assembly length, and the N50 and L50 values.

### Prediction of microbial composition, virulence factors and antimicrobial resistance

Profiling of the microbial community was performed on the processed reads using MetaPhlAn3.1 (v3.1.0, database “mpa_v31_CHOCOPhlAn_201901”) [60]. Simultaneously, profiling of antibiotic resistance factors was done using RGI (v5.2.0, CARD data v3.1.4, prevalence, resistomes and variants data v3.0.9) [61]. To obtain additional in-depth details of ARGs, in addition to the detection of virulence factors and mobile genetic elements (MGEs), PathoFact (v1.0; modified branch allowing the input of ORFs, #6fa64961) was run [62]. PathoFact is a pipeline for the prediction of antimicrobial resistance genes and virulence factors, and their localization to MGEs, in metagenomic data. PathoFact was run on the contigs assembled by IMP together with their predicted protein sequences (ORFs) for each sample separately. PathoFact uses DeepARG [63] and RGI [61] for the prediction of ARGs, DeepVirFinder [64] and VirSorter [65] for the prediction of phages and PlasFlow [66] for the prediction of plasmids. Additionally, PathoFact uses its own developed tool, a combination of a HMM database (built on the VFDB [67]) and a random forest model, for the prediction of virulence factors. To run PathoFact, the input protein sequences were first processed to remove any trailing stop codon symbols (“*”) and to remove any sequence having an internal stop codon symbol as this is required for the tool RGI for ARG detection.

### Data analysis

Statistical analysis of the taxonomic and functional data, as well as further visualisations, were done using version 4.1.1 of the R statistical software package [68]. The R package MaAsLin2 [69] was used to determine associations between the cohort data and microbial features (e.g., functional and taxonomic profiles). In addition, any identified significant differences were further validated by Wilcoxon rank-sum tests with adjustments using the ‘Benjamini-Hochberg’ method for multiple testing, specifically the ‘p.adjust’ function from the *stats* R package was used. The *tidyverse, microbiomeViz*, *tidytree* and *ggtree* packages were used to visualise the microbiome data, including the cladogram visualisation. The *tidyverse* package, including *ggplot2*, was used to generate all violin plots, box plots and bubble plots. Finally, the *hmisc* and *corrplot* packages were used for all correlation plots.

## Code and data availability

All code used for metagenomic and metatranscriptomic data analysis can be found in the following repository: https://gitlab.lcsb.uni.lu/ESB/co-infectomics. The repository at https://gitlab.lcsb.uni.lu/laura.denies/co-infectomics_r_analysis provides the code for the subsequent analyses in R. Individual workflows (sequencing data processing, QC, profiling, assembly, and analysis) were implemented using Snakemake [70]. Preprocessed and filtered reads, i.e., those filtered against the human genome (hg38), were submitted to the Sequence Read Archive hosted by NCBI and can be found under the accession PRJNA890008. Associated metadata is included in the code repository listed above. Additional data including assemblies, taxonomic, virome, functional pathway profiles along with MultiQC reports, can be found on Zenodo under the following link: https://doi.org/10.5281/zenodo.7192682.

## Supplementary figures

**Supplementary figure 1.**
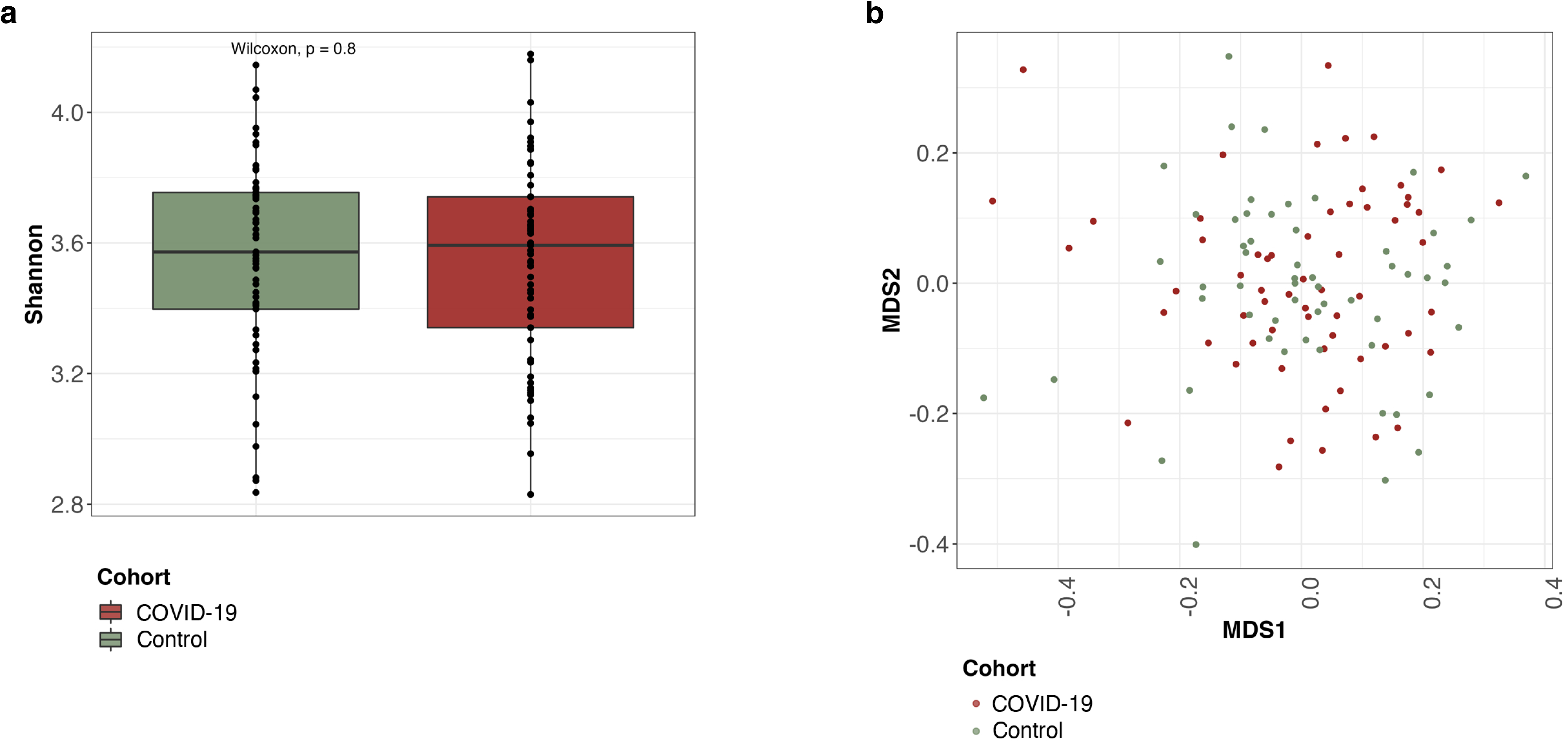
Dissimilarity of COVID-19 and control group microbiome profiles. Alpha-(Shannon) (**a**) and beta-diversity (**b**) metrics of the gut microbiome compositions of the COVID-19 and control groups.

